# Nucleation of the destruction complex on the centrosome accelerates degradation of β-catenin and regulates Wnt signal transmission

**DOI:** 10.1101/2022.02.01.478717

**Authors:** Ryan S. Lach, Chongxu Qiu, Erfan Zeyaei Kajbaf, Naomi Baxter, Dasol Han, Alex Wang, Hannah Lock, Orlando Chirikian, Beth Pruitt, Maxwell Z. Wilson

## Abstract

Wnt signal transduction is mediated by a protein assembly called the Destruction Complex (DC) made from scaffold proteins and kinases that are essential for transducing extracellular Wnt ligand concentrations to changes in nuclear β-catenin, the pathway’s transcriptional effector. Recently, DC scaffold proteins have been shown to undergo liquid-liquid phase separation *in vivo* and *in vitro* providing evidence for a mesoscale organization of the DC. However, the mesoscale organization of DC at endogenous expression levels and how that organization could play a role in β-catenin processing is unknown. Here we find that the native mesoscale structure is a dynamic biomolecular condensate nucleated by the centrosome. Through a combination of advanced microscopy, CRISPR-engineered custom fluorescent tags, finite element simulations, and optogenetic tools, that allow for independent manipulation of the biophysical parameters that drive condensate formation, we find that a function of DC nucleation by the centrosome is to drive efficient processing of β-catenin by co-localizing DC components to a single reaction hub. We demonstrate that simply increasing the concentration of a single DC kinase onto the centrosome controls β-catenin processing. This simple change in localization completely alters the fate of the Wnt-driven human embryonic stem cell differentiation to mesoderm. Our findings demonstrate the role of nucleators in dynamically controlling the activities of biomolecular condensates and suggest a tight integration between cell cycle progression and Wnt signal transduction.

## Introduction

The canonical Wnt signaling pathway is a highly-conserved^1^ morphogenic pathway that is essential for embryonic development, homeostatic maintenance of adult tissues and, when dysregulated, induces oncogenic malignancies^2–4^. Pathway signals converge onto a protein assembly called the destruction complex (DC) which tunes the stability of β-catenin (β-cat), the pathway’s central transcriptional effector, by regulating its interactions with the Wnt pathway kinases, CK1α and GSK3β, and the ubiquitinase, β-TRCP, which ultimately directs β-cat to the ubiquitin-mediated proteolysis machinery. Yet, despite the DC’s role in regulating β-cat stability, the structural principles that underly this protein assembly’s proper functioning in development and dysregulation in disease are still poorly understood.

In the Wnt OFF state, the DC maintains a low cytoplasmic concentration of β-cat by marking it, via phosphorylation and poly-ubiquitination, for proteasomal degradation. In the presence of extracellular Wnt ligand, Wnt/Frizzled/LRP5/6 complexes inhibit DC function through a mechanism that is still unclear, but likely involves selective recruitment of DC components to the signalosome, a liquid-like biomolecular condensate on the plasma membrane nucleated by Wnt/Frizzled/LRP5/6 clusters^5^. Indeed, optogenetic clustering of the cytoplasmic domain of LRP5/6 is sufficient to stabilize β-cat^6^. Thus, the formation of mesoscale protein clusters at the Wnt receptor level is necessary and sufficient for activating the pathway.

Aside from the core components and their protein-protein interactions^7^, less is known about the DC’s native structure, how that structure maintains low β-cat levels in the OFF state, and how structural changes to the DC might result in the accumulation of β-cat in the Wnt ON state. Recently, much interest has been paid to the propensity for the cytoplasmic Wnt pathway components to form mesoscale condensates. DC scaffolds Axin and Adenomatous Polyposis Coli (APC), as well as the signalosome ‘adapter’ protein Disheveled readily undergo liquid-liquid phase separation (LLPS) *in vitro* ^8^ and when exogenously over-expressed *in vivo*^9–11^. Cancer-causing mutations that eliminate Axin or APC LLPS cause aberrant accumulation of β-cat^7,12^ and can be rescued by fusing orthogonal protein-multimerizing domains^13^. These studies suggest that assembly of DC components into condensates is essential for normal DC function which begs the question: what is the role of mesoscale assembly of the cytoplasmic DC components in regulating β-cat stabilityã

One potential explanation is the “molecular crucible” model, which suggests that phase separation of DC components promotes efficient β-cat degradation in Wnt OFF conditions^8^. This theory posits that Axin and APC function as multivalent organizers that drive DC components to de-mix from the bulk of the cytoplasm, thereby concentrating the DC clients–CK1α, GSK3β, and β-cat— to effectively increase the rate of β-cat processing by locally concentrating pathway components. LLPS-driven partitioning of the DC is distinct from theories suggesting that the DC acts as a solid-state, assembly-line-like scaffold such as Ste5 that organizes the MAPK pathway in yeast^14^. In support of this theory, small deletions in Axin’s Intrinsically Disordered Region (IDR), which drive LLPS^15^, increase β-cat stability and downstream gene activation^8^. In this paradigm, conditions that alter the phase behavior of scaffolds and partitioning coefficient of clients are predicted to regulate the stability of β-cat and Wnt pathway signal transmission.

One potential mechanism for regulating the time and place where condensates form is to control nucleation sites of intracellular droplets. This biophysical mechanism was recently explored *in vivo*, using a synthetic optogenetic system^16^. Yet the role of nucleation in regulating the function of naturally-occurring biomolecular condensates remains unexplored. Since phase separating systems exhibit switch-like responses to changes in concentration^17,18^ and can have biophysical parameters tuned to exist near these transitions *in vivo*^19,20^, dissecting the natural regulatory mechanisms is difficult without fine control over protein concentration and affinity. This suggests that overexpression of DC scaffolds may not accurately reflect its endogenous biophysical state and mesoscale structure, which could lead to misinterpretation of the role of LLPS of DC scaffolds in regulating β-cat stability. Here we overcome this barrier by utilizing CRISPR Cas9 gene editing, custom inducible expression vectors, and optogenetic tools to observe and probe the native, mesoscale organization of the DC in the Wnt OFF, Wnt ON states.

Building on results demonstrating that β-cat^21^, Axin1^22^ and APC^23^ localized to the centrosome, we show that all DC components are nucleated by the centrosome into dynamic, liquid-like biomolecular condensates. In support of the molecular crucible theory, we find that nucleation drives efficient degradation of β-cat. We demonstrate that centrosomal nucleation is controlled by Axin1, but not APC concentration, suggesting the existence of an intracellular phase boundary that the DC crosses via Axin1 concentration. We then utilize a Cahn-Hilliard-based simulation of DC droplet formation and enzyme kinetics to predict how nucleation and affinity of DC components promotes efficient β-cat processing. Finally, using our model as a guide we engineered a light-inducible GSK3β (Opto-GSK3) to control its partitioning at the centrosome, which increases β-cat degradation as predicted by our computational model. Opto-GSK3 activation is so effective that it inhibits Wnt-driven differentiation of human stem cells into mesoderm. These findings show that DC droplet formation is nucleated by the centrosome and suggest that DC scaffolds function to concentrate clients in liquid droplets *in vivo* to accelerate the degradation of β-cat.

## Results

### β-catenin condensation is predictive of Wnt pathway activity state

To understand the role of mesoscale organization in DC function, we first sought to characterize the DC’s main substrate, β-cat, in single, live cells. We used CRISPR-Cas9 to knock-in a custom fluorescent tag, tdmRuby3, to the N-terminus of the *CTNNB1* gene of 293T cells (**Fig. 1A**). Live-cell confocal imaging revealed the expected cytoplasmic accumulation in response to Wnt-3a ligand and the GSK3β inhibitor CHIR (**Supp. Fig. 1A-B, Supp. Vid. 1**) and expected localization of β-cat at the cell membrane with enrichment at cell-cell contacts, consistent with previous work in fixed specimens^24^. In addition to these commonly observed pools, we observed nearly every cell contained 1-2 bright, spherical, perinuclear β-cat puncta (**Fig. 1B** *top left*). Timelapse imaging showed fission and fusion of puncta on the timescale of minutes (**Supp Fig. 1C)**, suggesting that these structures are liquid-like biomolecular condensates. Given the prevalence of biomolecular condensates in organizing processes in mammalian cell biology we hypothesized that these perinuclear puncta might organize β-cat destruction.

**Fig. 1:**
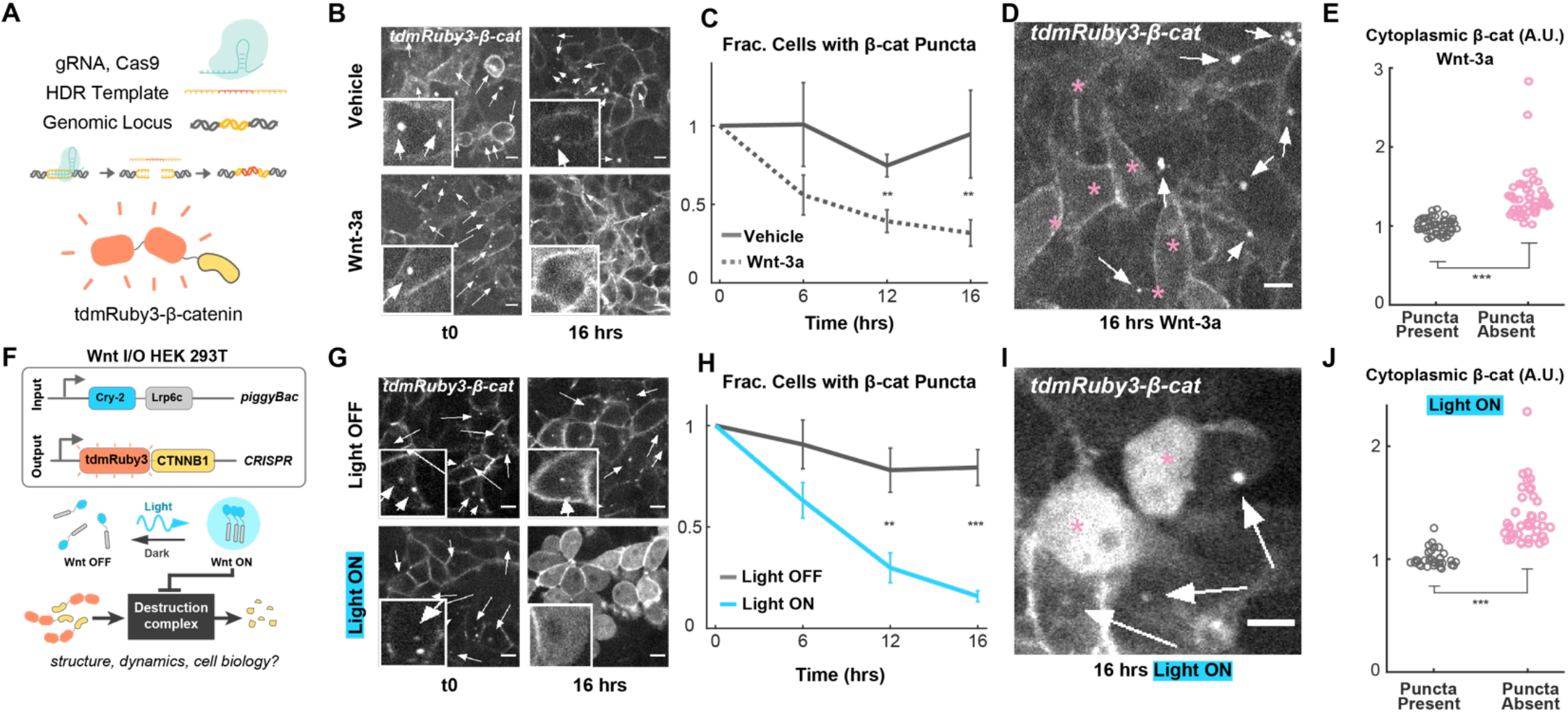
Endogenously expressed β-catenin puncta are inversely correlated with lrp6-mediated Wnt pathway activation and β-catenin accumulation. **A**. Schematic of tdmRuby3 CRISPR tag strategy. **B**. Representative tdmRuby3- β-catenin images of cells treated with Wnt-3a or media vehicle. Arrows indicate β-catenin puncta. Scale = 10μm. **C**. Fraction of t0 population with visible β-catenin puncta, presented as mean +/- s.e.m. (N=12 imaging fields per condition). **D**. Representative cells from Wnt-3a condition. Arrows indicate puncta, asterisks indicate cells lacking puncta. **E**. Comparison of mean cytoplasmic β-catenin fluorescence between Wnt-3a cells with and without visible β-catenin puncta. **F**. Schematic of Wnt I/O cells containing lentivirally-expressed Cry2-LRP6c and CRISPR-tagged tdmRuby3-β-catenin. Stimulation of Cry-2-Lrp6c with blue light results in reversible clustering of lrp6c and downstream pathway activation. **G**. Representative tdmRuby3-β-catenin images of cells stimulated with blue light or left in the dark throughout imaging timecourse. **H**. Fraction of t0 population with visible β-catenin puncta, presented as mean +/- s.e.m. (N=12 imaging fields per condition). **I**. Representative cells from Wnt-3a condition. Arrows indicate puncta, asterisks indicate cells lacking puncta. **J**. Comparison of mean cytoplasmic β-catenin fluorescence between Light ON cells with and without visible β- catenin puncta.

To determine if Wnt pathway activation altered the perinuclear puncta we performed volumetric confocal timelapse microscopy on our *tdmRuby3-β-cat* cells and quantified the fraction of cells with puncta as a function of Wnt-3a ligand treatment and time. At the population level, the fraction of cells with puncta significantly decreased in response to Wnt-3a (**Fig. 1C**). We found this same relationship existed between single cells in an isogenic population, with non-responding cells maintaining their puncta and responding cells dissolving them (**Fig. 1D-E**). Thus, the disappearance of perinuclear β-cat puncta is correlated with β-cat accumulation and the existence of these puncta is correlated to the resistance of ligand-induced accumulation.

To establish if directly activating the Wnt receptor controls the existence of the puncta we transduced *tdmRuby3-β-cat* cells with an optogenetic version of the Wnt co-receptor, LRP6c (Opto-LRP6)^6^. Opto-LRP6 induced greater accumulation of β-cat than either Wnt or CHIR (**Supp Fig. 1D**). We thus reasoned that this all-optical Wnt input control and output visualization cell line would maximize our ability to observe rearrangements in pathway components due to a higher dynamic range of activation (**Fig. 1F**). We found that activating Opto-LRP6 resulted in an even greater reduction in the fraction of cells containing β-cat puncta than ligand treated cells (**Fig. 1G-H, Supp. Vid. 2-3**). Further, of light-stimulated cells, those that were resistant to optogenetic activation maintained their β-cat puncta (**Fig.1 I-J**). We also observed this same resistance to β-cat accumulation in response to CHIR (**Supp Fig. 1E**). Together these results indicate that activation of the Wnt pathway causes perinuclear puncta to dissolve and the presence of these puncta is inversely related to Wnt pathway activation at both the population and single-cell level.

### The destruction complex forms a biomolecular condensate colocalized to the centrosome

We next sought to determine (i) what, if any, cellular structure was organizing these puncta, (ii) if all DC components were co-localized in the puncta, and (iii) whether these were solid-or liquid-like condensates. Because of the sensitivity of LLPS systems to protein concentration^25^ we decided on a strategy that allowed us to visualize DC components at low or endogenous concentrations, while retaining the ability to assess protein dynamics through live-cell microscopy and Fluorescence Recovery After Photobleaching (FRAP). Indeed, the DC scaffolds (APC and Axin1) form multiple cytoplasmic liquid droplets when overexpressed^12,26^. Thus, we used CRISPR to knock-in tdmRuby3 into the loci of *CSNK1A1* and *GSK3B*, the genes encoding the kinases CK1α and GSK3β that sequentially phosphorylate β-cat in the DC.

We found that both DC kinases were localized into 1-2 perinuclear puncta (**Fig. 2A**). Timelapse imaging revealed that the number and position of CK1α and GSK3β puncta were determined by cell cycle stage (**Supp. Fig. 2A**): we observed single condensates in G1, two condensates in G2/S, and a ‘finger’-like pattern—suggesting association with the mitotic spindle— during late mitosis. These observations, combined with previous reports of perinuclear enrichment of CK1α and GSK3β in fixed cells^27,28^, led us to hypothesize that both kinases and β-cat were associated with the centrosome. Immunofluorescence staining for γ-tubulin (**Fig. 2B**) and GM130 (**Supp. Fig. 2B)**, canonical centrosome markers, confirmed that tdmRuby3-CK1α, tdmRuby3-GSK3β and tdmRuby3-β-cat puncta were indeed colocalized to the centrosome.

**Fig. 2:**
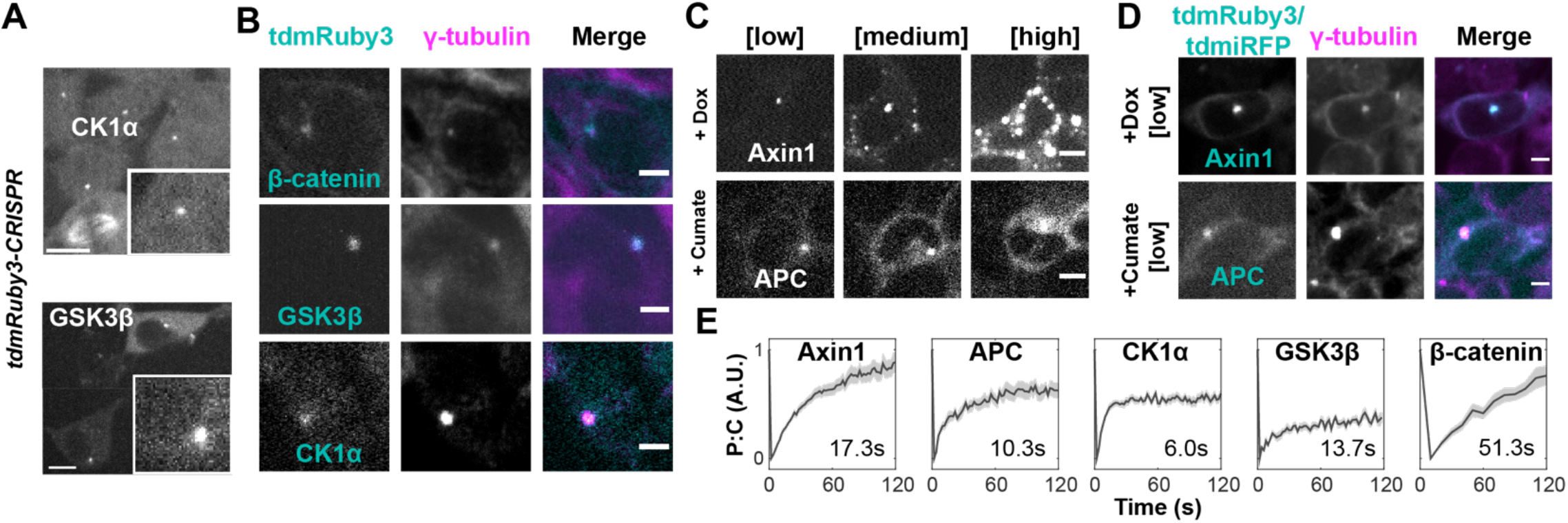
Canonical Destruction Complex (DC) components reside in liquid droplets nucleated at the centrosome. **A**. Representative images of CRISPR-integrated tdmRuby3-CK1α and tdmRuby3-GSK3β cells. *Insets*: closeup views of singular peri-nuclear puncta. Scale = 10μm. **B**. Representative cells bearing the indicated DC component fixed and stained for endogenous γ-tubulin. Scale = 10μm. **C**.Representative timelapse images from live cells bearing dox- and cumate-inducible Axin1 and APC cassettes under induction. Montages depict the same cell increasing its DC scaffold concentration through time. **D**. Representative cells bearing the indicated DC component fixed and stained for endogenous γ-tubulin. **E**. Fluorescence recovery after photobleaching (FRAP) traces of mean puncta:cytoplasm fluorescence ratio for indicated DC components. Data presented as mean +/- s.e.m. normalized to extent of bleaching (N=39, 20, 33, 17, 22 for Axin1, APC, CK1α, GSK3β, β-catenin respectively). Individual FRAP traces were fit to the equation: f(t) = a(1-e^(-bt)^) to obtain a and b parameters and half-max recovery time (τ1/2). Mean τ1/2 for each DC component is displayed on each plot.

Localization of the DC kinases to the centrosome has not been associated with β-cat degradation. Thus, we wondered if the DC scaffolds Axin1 and APC, which are required for efficient processing of β-cat, are also localized to the centrosome. To test whether Axin1 and APC are localized to the centrosome at low cellular concentrations, but not when overexpressed, we generated clonal 293Ts bearing doxycycline (Dox)-inducible human *Axin1-tdmRuby3* and cumate-inducible human *APC-tdmiRFP670*. At low levels of induction, both Axin1 and APC localization mirrored CK1α, GSK3β and β-cat, forming bright perinuclear puncta (**Fig. 2C** *left*) that colocalized with centrosomal markers (**Fig. 2D, Supp. Fig. 2B)** and replicated following cell cycle progression (**Supp. Fig. 2A**). As protein concentration increased, Axin1 but not APC formed extra-centrosomal puncta throughout the cytoplasm (**Fig. 2C-E**). 293Ts are commonly used in experiments probing DC mesoscale structure *in vivo*^8,11^, but expression of Wnt pathway components may vary significantly between stem cells and differentiated cells. We observed the same preferential localization of Axin1 at low concentration in human induced-pluripotent stem cells (iPSCs) (**Supp. Fig. 2F)**. These findings establish that all DC components necessary for phosphorylating β-cat, prior to its ubiquitination, are localized at the centrosome throughout the cell cycle and suggest that DC centrosomal nucleation is generalizable to multiple cell types.

When overexpressed Axin and APC cross the phase boundary and form liquid condensates in the cytoplasm that are hypothesized to concentrate DC kinases and β-cat, thereby organizing β-cat degradation^29^. The fact that no extra-centrosomal DC-puncta were observed in cells at endogenous concentrations led us to hypothesize that the DC is a liquid organelle that is nucleated at the centrosome at endogenous protein concentrations. Next, we sought to determine the material state of the centrosomal DC using FRAP on CRISPR-tagged CK1α, GSK3β and β-cat, as well as on Axin1 and APC at low levels of induction. All centrosomal DC components exhibited mean half-maximal recovery times (τ1/2) between 10 and 60s (**Fig. 2E**)—like in over-expressed systems^15^ and in-line with mesoscale cellular structures classically considered liquid-like^30^. These results indicate that the DC is a liquid nucleated by the centrosome and suggest that nucleation has a role in maintenance of cellular β-cat levels.

### A reactive Cahn-Hilliard model predicts accelerated b-catenin processing upon centrosomal nucleation of DC clients

To understand the effect of centrosomal nucleation of DC components on β-cat processing we simulated the processive phosphorylation of β-cat by DC kinases, using a reactive, multi-component, Cahn-Hilliard system^31,32^. We represented the function of DC scaffolds implicitly through the interaction parameters between kinases and β-cat **(Fig. 3A)**. Indeed, synthetic DC scaffolds with these simple attributes have been shown to rescue aberrant Wnt signaling^33^.

**Fig. 3:**
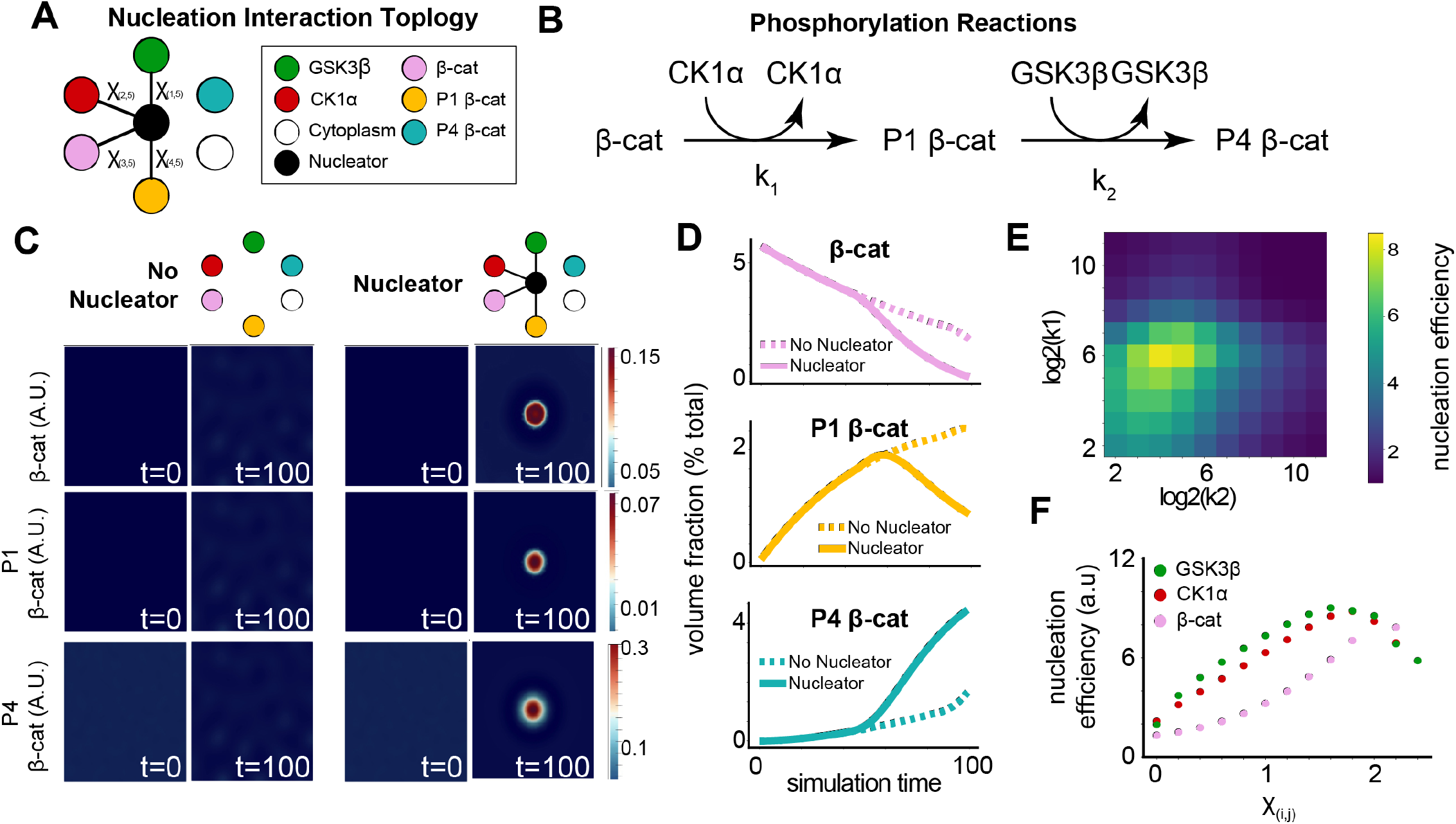
*In silico* modelling of β-catenins processing efficiency from a nucleated liquid droplet. **A**. Nucleation interaction topology that describes the interactions between each component of the simulation. Connected components minimize free energy by mixing and unconnected components either de-mix or remain in a non-interacting neutral state. **B**. Schema describing the phosphorylation reactions and rates modeled in the simulation. **C**. Simulation at steps 0 and 100 comparing a system with and without a centrosome. **D**. Quantification of each form of β-catenin with and without a centrosome. **E**. Nucleation efficiency as a function of both rate parameters k1 and k2. **F**. Nucleation efficiency in simulations as a function of the interaction parameters between a single client and the cytoplasm.

In our simulation, we represent the volume fraction of each DC protein *ϕ*_*i*_, as an incompressible volume such that 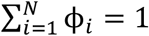 and approximate the reaction rates with spatially dependent analogues to well-mixed reactions using the simplified, non-state dependent description of the second order rate *R*_*i*_ = *k*_*i,j*_ *ϕ*_*i*_ *ϕ*_*j*_, with production and consumption denoted by the sign of *k*_*i,j*_^34,35^ (**Fig 3B**). Thus, our reactive Cahn-Hilliard simulation is modeled as two coupled second order equations

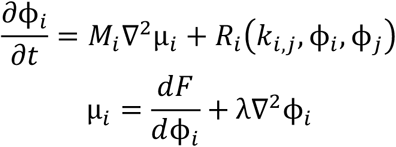

Where *M*_*i*_ is the mobility constant, with all DC components having the same diffusion rate, λ, is the surface energy parameter that dictates the length of transition regions between domains, and *F* is the polynomial double-well description of the free energy:

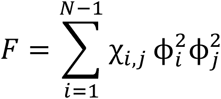

Where, χ_*i,j*_, describes interaction strength between DC proteins, the cytoplasm, and the centrosome. We modeled centrosomal nucleation as a region in the simulation with increased interaction strength as has been done previously to describe nucleation sites^36^ (**Supp. Fig. 3A**). To determine the size of this nucleation region we measured the relative volume of centrosomally-localized DC kinases and β-cat (**Supp. Fig. 3B**).

To test the effects of nucleation on β-cat processing we compared simulations in the presence and absence of a nucleation region (**Fig. 3C**). We found that for systems that did not spontaneously phase separate, which mimics the endogenously expressed conditions observed above, DC components localized into a single droplet surrounding the nucleator but do not spontaneously de-mix in its absence**(Sup Video 6, 7)**. We found that the nucleated system processed β-cat and its intermediates more quickly (**Fig. 3D**) over a wide range of nucleator sizes (**Supp Fig. 3E**). Notably, the nucleated system accelerated β-cat processing, increasing pathway efficiency (**Supp. Fig. 3F**). This efficiency gain was maintained over a large range of reaction rates (**Fig. 3E, Supp Fig. 3G**). As expected, in systems with high reaction rates–three orders of magnitude greater than the mobility constant–the effect of nucleated phase separation is no longer observed.

Given our findings that nucleation drives efficient processing of β-cat we hypothesized that χ, the interaction parameter that drives phase separation, is a control parameter for β-cat processing. To determine the relationship between DC function and the interaction strength parameter we systematically decreased the χ between DC clients and the cytoplasm. We found that driving less condensation on the nucleator, through altering χ, decreased the speed and efficiency of β-cat processing (**Supp Fig 3H-I, Fig. 3F**). Together these results demonstrate that nucleation of DC components has the potential to increase β-cat processing and that a tunable control parameter of this process is the free energy of mixing.

### Optogenetically-driven Enrichment of Centrosomal GSK3β Condensates Rescues Hyperactivated Wnt signaling

*In silico* analysis of the DC suggests that processing efficiency in the presence of a nucleator dependent on client condensation. Imaging of GSK3β showed relatively weak enrichment in centrosomal puncta compared to CK1α, leading to the idea that increasing nucleation of GSK3β will increase the degradation rate of β-cat *in vivo*. Changing concentration alters both the system’s propensity to undergo spontaneous LLPS and its reaction rates^37^ and therefore cannot be used to test the effect of nucleation on a reaction. On the other hand, optogenetic photo-clustering domains can independently control intracellular LLPS at fixed concentrations via light-induced changes in valency between monomers ^38,39^. Thus, we reasoned that an optogenetic tool that drives changes in free energy could isolate the effect of phase from biological function.

To test if photo-clustering could increase partitioning to a nucleator, we fused the photo-oligomerizer Cryptochrome-2 (Cry-2) and eGFP to the N-terminus of the human GSK3β ORF (‘Opto-GSK3’ hereafter) and stably transduced this construct into 293Ts (**Fig. 4A**). Upon light stimulation, Opto-GSK3 increased its centrosomal enrichment, doubling the mean centrosome:cytoplasm fluorescence ratio within 10 seconds of activation (**Fig. 4B**,**C, Supp Vid. 7**). Notably, activation of Opto-GSK3 strictly resulted in the formation of 1 or 2 perinuclear puncta and did not form extra-centrosomal condensates, in contrast to studies using Cry2 alone^17^. Thus, we found that illumination of opto-GSK induced condensate formation only at the centrosome.

**Fig. 4:**
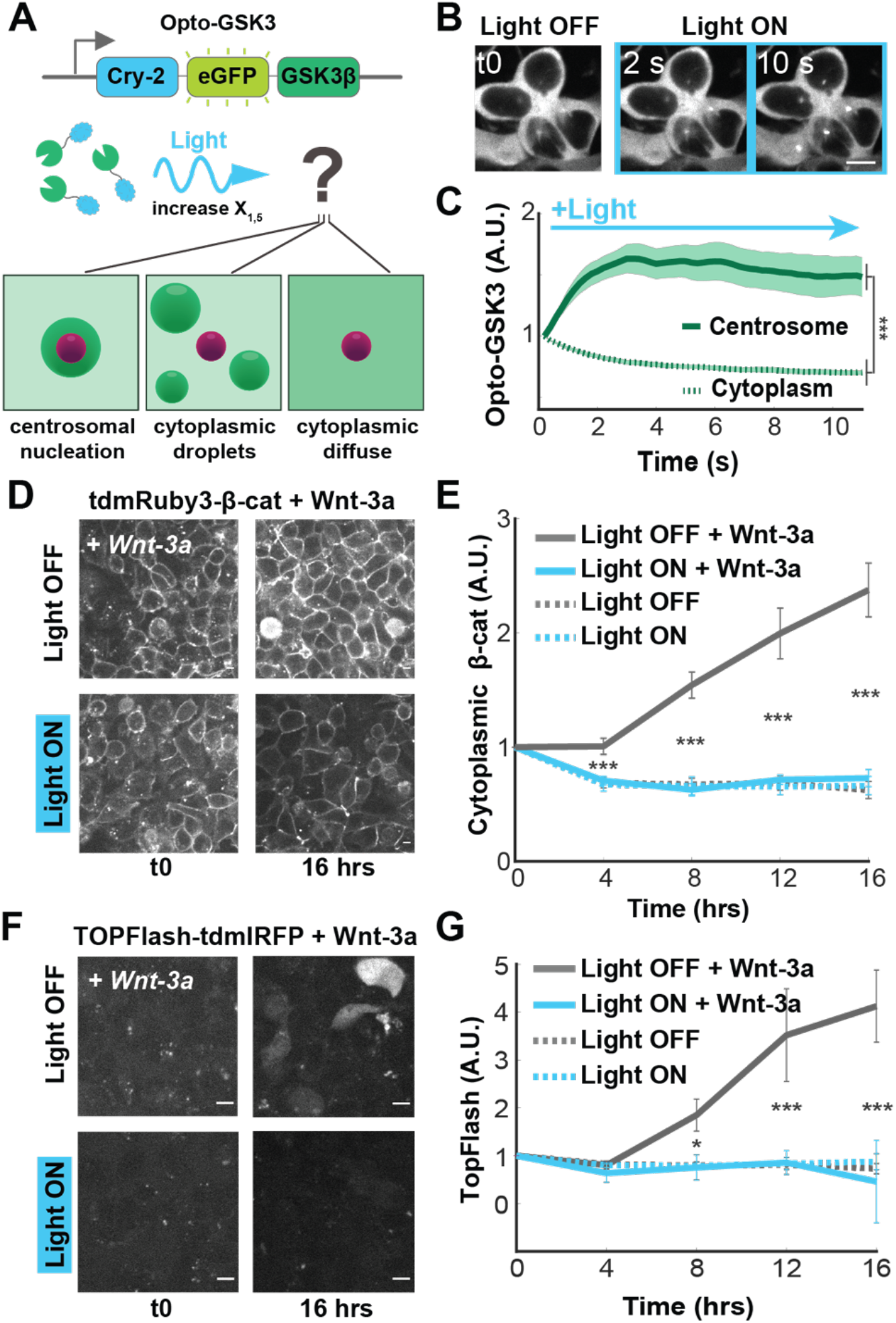
Optogenetic clustering of GSK3β increases centrosomal droplet partitioning and suppresses Wnt pathway activation. **A**. Schematic of Opto-GSK3 and possible spatial outcomes of blue light stimulation. **B**. Representative images of cells bearing Opto-GSK3 responding to blue light stimulation. Montage depicts the same cells throughout the activation timecourse. Scale = 10μm. **C**. Quantification of cells in **B:** Mean fluorescence fold-change from t0 for each compartment +/- s.e.m. (N=20 cells). **D**. Representative images of cells bearing Opto-GSK3 + tdmRuby3-β-catenin following treatment with Wnt-3a. Scale = 10μm. **E**. Quantification of cells in **D:** Mean fluorescence fold-change from t0 +/- s.e.m. is shown (N=20 cells per condition). **F**. Representative images of cells bearing Opto-GSK3 + TOPFlash-IRFP following treatment with Wnt-3a. Scale = 10μm. **G**. Quantification of **F:** Mean fluorescence fold-change from t0 +/- s.e.m. is shown (N=24 cells per condition).

To determine whether increased centrosomal condensation of GSK3β controlled Wnt signal transmission we activated Opto-GSK3 in cell lines with three distinct methods for increasing the cellular concentration of β-cat: ligand-induced, kinase inhibition, and dox-induced gene upregulation. We found that opto-GSK activation abolished both Wnt-3a-induced β-cat accumulation and transcriptional activation as measured by TOPFlash fluorescence (**Fig. 4D-E**). Control experiments comparing cells in light vs. dark confirmed that this was not due to light alone (**Supp. Fig. 4A-B**). We observed a similar effect when analyzing total β-cat by Western blotting and immunofluorescence staining (**Supp Fig 4C-E**). Given the modest accumulation of β-cat in response to Wnt-3a in 293Ts we tested to see if Opto-GSK3 clustering was sufficient to blunt β-cat accumulation induced by either CHIR or from a Dox-inducible β-cat over-expression construct. Indeed, activation of Opto-GSK3 also inhibited both methods for driving b-catenin accumulation in a light-dependent manner (**Supp Fig F-J**). These results demonstrate that increasing DC client nucleation at the centrosome can be used to dictate Wnt signal transmission across a wide range of activation regimes.

### Centrosomal Enrichment of GSK3β Prevents Wnt Pathway Activation-Induced Differentiation of Embryonic Stem Cells

Changes in β-cat concentration differentiate a variety of stem cell populations, including human embryonic stem cells (hESCs)^38,40^. Having determined that increased centrosomal nucleation of GSK3β is sufficient to reduce β-cat accumulation and Wnt-responsive gene transcription in 293T cells, we wondered whether it was also sufficient to prevent the downstream differentiation of hESCs. Both CHIR and Wnt-3a induce hESC differentiation into mesoderm which is monitored by Brachyury (BRA) expression^41^. To test whether centrosomal nucleation prevents differentiation, we expressed Opto-GSK3 in H9 hESCs and treated them with CHIR or DMSO control in the presence or absence of activating blue light for 24 hrs. Following stimulation, cells were fixed and stained for BRA to assay for differentiation. In the dark, hESCs receiving CHIR responded robustly, displaying bright nuclear BRA compared to DMSO controls (**Fig. 5A** *left*). However, when stimulated with blue light, CHIR-treated cells showed significantly reduced levels of BRA staining compared to the dark controls, indicating that nucleation of GSK3b countered CHIR-induced differentiation into mesoderm (**Fig. 5A** *right*). Interestingly, we observed that BRA levels in light-treated cells, in the DMSO control condition were slightly, but significantly higher than when in the dark, suggesting that over-repression of the Wnt pathway by Opto-GSK3 activation weakly promotes differentiation in hESCs as well (**Fig. 5B**).

**Fig. 5:**
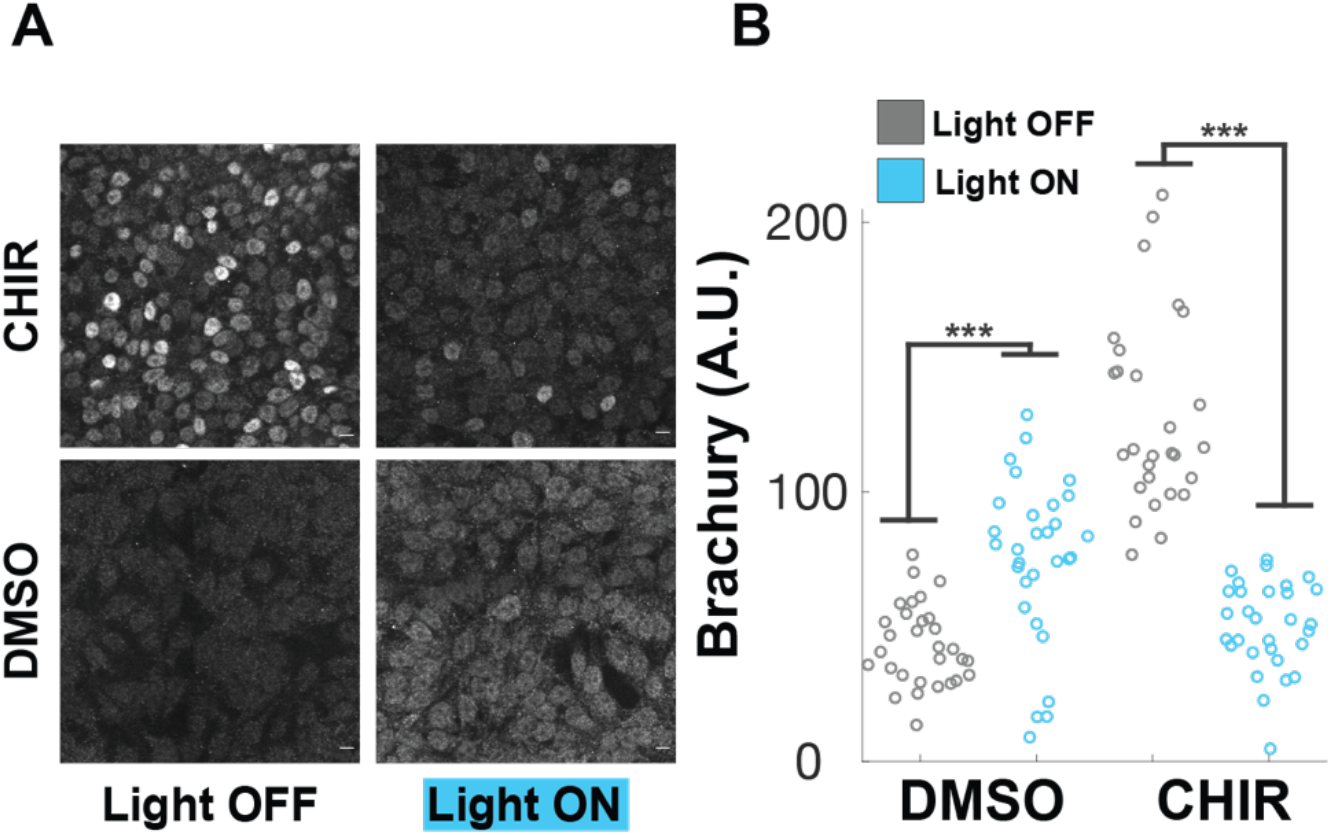
Optogenetic clustering of GSK3β suppresses Wnt pathway-mediated differentiation of embryonic stem cells. **A**. Representative images of H9 embryonic stem cells bearing Opto-GSK3 following 24hrs in described conditions, fixed and stained for endogenous Brachyury. **B**. Quantification of experiment from **A:** Mean nuclear fluorescence for cells measured in each condition is presented.

## Discussion

Canonical Wnt signaling has been the subject of intense interest for its importance in developmental decisions and dysregulation in a variety of cancers. Many studies have attempted to elucidate the role of the DC in regulating Wnt signal transduction, and with recent discoveries suggesting that LLPS plays a role in DC structure, we sought to understand how the biophysics of DC proteins regulate DC function in live, single cells. Through a combination of super-resolution microscopy, *in silico* modeling, and optogenetic methods to isolate and probe the phase diagram, we uncover the mesoscale structure of the DC to be a liquid condensate nucleated by the centrosome. The complementarity of these methods allowed us to identify a function for nucleation: it accelerates the catalytic action of DC proteins thereby promoting efficient processing of β-cat. Differential partitioning of DC clients in centrosomal condensates constitutes a novel biophysical parameter that controls Wnt signal transmission and suggests another potential function of coordinating Wnt signals with cell cycle control.

LLPS has been correlated with DC function *in vivo*^7^ and *in vitro*^8^. Yet, because of the sensitivity of protein phase separation to concentration we sought to examine the biophysics of DC compoents at their endogenous concentrations. To achieve this, we used custom fluorescent reporters to CRISPR the endogenous loci of β-cat and the two DC kinases, CK1α and GSK3β. To understand the control parameters of the DC structure we developed finely tunable, inducible expression vectors of the two scaffolds, APC and Axin1. The presence of many cytoplasmic Axin1 and APC droplets in mildly (approx. 4X) overexpressed cellular conditions^10^ has been cited in support of the idea that DC scaffolds spontaneously demix at endogenous concentrations. We find evidence against this in human 293T, human iPSC and hESCs. In contrast, we found that at low or endogenous levels all DC components form dynamic assemblies with preferred localization to the centrosome, highlighting the importance of studying phase separating systems *in vivo* at endogenous concentrations.

Our results support a ‘molecular crucible’ model of β-cat degradation, in which DC scaffolds serve to concentrate DC clients in nucleated droplets to increase β-cat phosphorylation rate. Nanoscale models for basal β-cat degradation have been proposed, including a radial Axin structure that complexes with a ‘ratcheting’ APC to sequester, sequentially phosphorylate and release β-cat to associate with βTrCP for ubiquitination^15^, an APC-mediated regulatory loop that tunes phosphorylation rate via dynamic degradation of Axin^42^.The present findings point toward LLPS as a mechanism to concentrate sparse clients in the cytoplasm. *In vitro* Axin1 polymerization has been observed to be ordered^26^,yet others have shown that Axin1 and APC form surface tension-minimizing spherical droplets that fuse over time^43^ whose existence is determined by concentration and interaction strength^10^. Our results suggest that nucleation to the centrosome is central to the formation of this molecular crucible and future studies examining the modulation of DC component properties at the centrosome promise to yield new steady-state characteristics and regulatory mechanisms of Wnt signaling dynamics.

Our results raise an important question that may lead to the discovery of unknown potentiators of Wnt signal transduction. What is/are the nucleator(s) coupling the DC to the centrosomeã Axin1 is known to associate with γ-tubulin^44^ and is a substrate of PLK1, a kinase involved in centrosome duplication during cell cycle progression^22^, suggesting that it is tightly associated with the centrosome. APC is a regulator of microtubule stability and growth^23,45^ and its armadillo repeat region—which also binds Axin^33^—is sufficient to induce centrosomal localization^23^. The high degree of multivalency and inter-DC-component interactions suggest that multiple interactions could localize the DC to the centrosome, increasing the robustness of nucleation. Proximity labeling experiments promise to help elucidate the potential interactions.

Finally, centrosomal nucleation of the DC suggests a potential function in coordinating cell cycle progression with Wnt signaling. We found two DC droplets in cells with duplicated centrosomes. This suggests an efficient method to partition DC components during mitosis. Non-nucleated droplets would be subject to random segregation to daughter cells, leading to potentially detrimental asymmetry in subsequent Wnt signaling capacity of the growing tissue. Thus, cell cycle synchronization could be a method of reducing heterogeneity in Wnt-induced stem cell differentiations. Additionally, as the main microtubule-organizing center in the cell, it is possible that the centrosome integrates Wnt signal transduction with cytoskeletal state to align Wnt-driven cell fates with mechanical cues.

Overall, our studies suggest an integral role for nucleation of LLPS in regulating the activity of membraneless organelles *in vivo*. The power of observing proteins in their endogenous contexts coupled to the ability to precisely tune interaction strength, without altering protein function or concentration, enables the functional dissection of membraneless organelles.

## Acknowledgements

We thank Ken Kosik and Denise Montell for guidance on the project direction, Cassidy Arnold and Erik Hopkins for technical support with maintenance, enrichment and assaying clonal cell lines, and Imogen Rawlings-Green, Natalie Tjahjadi and Mark Lu for support with data analysis. We thank R.L. the Eunice Kennedy Shriver National Institute of Child Health and Development for funding R.L. (1F31HD106900-01A1) and CRCC 2021-22 Faculty Seed Grant (UCSB, C22CR4129) for funding the work.

## Materials and Methods

### Cell Lines

Human 293T cells were cultured at 37°C and 5% CO2 Dulbecco’s Modified Eagle Medium, high glucose GlutaMAX (Thermo Fisher Scientific, 10566016) medium supplemented with 10% fetal bovine serum (Atlas Biologicals, F-0500-D) and 1% penicillin-streptomycin. Human induced pluripotent stem cell line (hiPSC) WTC was gifted by the Pruitt lab (purchased from Coriell). hiPSCs were propagated on Matrigel® coated tissue culture plates using serum-free essential 8 (Gibco) culture conditions in standard environments consisting of 5% carbon dioxide at 37°C. Experiments in human Embryonic Stem Cell (hESC) lines were performed using the H9 hESC cell line purchased from the William K. Bowes Center for Stem Cell Biology and Engineering at UCSB. Cells were grown in mTeSR™ Plus medium (Stem Cell Technologies) on Matrigel® (Corning) coated tissue culture dishes and tested for mycoplasma in 2-month intervals.

### Cloning of PiggyBac Transposase and Lentiviral Overexpression Constructs

#### pPig_H2B-mTagBFP2::t2A::Cas9-Avidin

was constructed via subcloning human H2B, mTagBFP2 and Cas9-Avidin provided by Max Wilson into an expression vector bearing a CMV promoter and flanking PiggyBac transposase-compatible inverted terminal repeats using Gibson Assembly (New England BioLabs Inc., E2611L) according to supplier instructions. Each of the PCR fragments used were amplified using the following primers:

PiggyBac (CMV) Backbone fwd: tgacgcccgccccac rev: ggtaagctttttgcaaaagcctaggcc. H2B + 18AA linker fwd: cctaggcttttgcaaaaagcttaccatgccagagccagcgaagtc rev: GCATATTTTCCTTGATGAGTTCACTCATccCagTatGtcCgcCggAg. mTagBFP2 fwd: ATGAGTGAACTCATCAAGGAAAATATGCACATG rev: CGTCCCCGCAGGTCAACAAACTTCCGCGACCTTCTCCGCTCCCATTGAGCTTATGGCCGAGTTT GCTG. 3X-Flag-NLS-Cas9-HA-Avidin fwd: GGAAGTTTGTTGACCTGCGGGGACGTGGAAGAAAACCCGGGTCCAgactataaggaccacgacggagact ac rev: gctgcgggtcgtggggcgggcgtcaggatccagacgccgcag

#### XLone-Axin-tdmRuby3

was constructed via PCR and Gibson Assembly, subcloning from the following constructs: Flag-Axin1 purchased from Addgene (#109370), tdmRuby3 from Max Wilson into XLone-GFP purchased from Addgene (#96930) containing 3^rd^ gen tet ON-responsive promoter and EF1α-driven Blasticidin selection cassette. The following primers were used: XLone Backbone fwd: taaactagtagaccacctcccctgcg, rev: ggtacctttacgagggtaggaagtgg, human Axin1 fwd: cacttcctaccctcgtaaaggtaccatgaatatccaagagcagggtttcccc, rev: CCATgctTCCgCCgCCACTACCgCCgtccaccttctccactttgccgatgatc, 7AA link-tdmRuby3 fwd: GGcGGTAGTGGcGGcGGAagcATGGTTAGCAAAGGGGAGGAGC, rev: gcaggggaggtggtctactagtttaCTTGTACAGCTCGTCCATGCCG.

#### XLone-bCat-tdmRuby3

was constructed via PCR and Gibson Assembly, subcloning from the following constructs: XLone-Axin-tdmRuby3 (above) and Human Beta-catenin GFP purchased from Addgene (#71367). The following primers were used: XLone Backbone fwd: GGcGGTAGTGGcGGcGGAagcATGGTTAGCAAAGGGGAGGAGC, rev:

ggtacctttacgagggtaggaagtgg, human bcat fwd: cacttcctaccctcgtaaaggtaccatggctactcaagctgatttgatggagttg, rev:

CCATgctTCCgCCgCCACTACCgCCcaggtcagtatcaaaccaggccagc

#### pPig_CuO-APC-tdmIRFP670

CymR was constructed via PCR and Gibson Assembly from the following constructs: pCuo CA Rac1 CMV + cumate operon purchased from Addgene (#84643), human APC open reading frame purchased from Addgene (#16507), tdmirfp670nano from Max Wilson, human ubiquitin C-driven CymR Cuo repressor purchased from Addgene (#119907) into pPig-Hygro transposase backbone from Max Wilson. PCR fragments were amplified using the following primers:

pPig-Hygro Backbone fwd: GGACGTGGAAGAAAACCCGGGTCCAatgggtaaaaagcctgaactcaccgc, rev: cattccacagggtcgacagtacaagc,

Cuo + CMV fwd: cttgtactgtcgaccctgtggaatgcgttacataacttacggtaaatggcccgc, rev: actgatcatatgaagctgcagccatgaattcggtaccggatccagtcgactag,

APC fwd: atggctgcagcttcatatgatcagttgttaaagcaag,

rev: CCATgctTCCgCCgCCACTACCgCCaacagatgtcacaaggtaagacccagaatg, 7AAlinker-tdmirfp670nano fwd: GGcGGTAGTGGcGGc,

rev: ggcgccaaaacccggcgcggaggccttaGGACTGCTGTATTGCAATGCCAACTAC, UbC-CymR-V5-T2A fwd: ggcctccgcgccggg,

rev: TGGACCCGGGTTTTCTTCCACGTCCCCGCAGGTCAACAAACTTCCGCGACCTTCTCCGCTCCCc gtagaatcgagaccgaggagagg

#### pLV_Cry2-tdeGFP-GSK3b

was obtained via synthesis and cloning services provided by Vector Builder Inc. Full details available upon request, but briefly: primary plasmids containing *Arabadopsis thaliana*, tdmIRFP from Max Wilson and human GSK3β purchased from Addgene (# 16260) ORFs were supplied to VectorBuilder for cloning and EF1α-driven expression into 3^rd^ generation lentiviral backbone. Vectorbuilder provided the desired final, sequenced plasmid.

#### pPig_8XTOP_tdIRFP_Puro

was constructed via PCR and Gibson Assembly from the following constructs: pPig_H2B-mTagBFP2::t2A::Cas9-Avidin (above), M50 Super 8x TOPFlash purchased from Addgene (#12456) and codon-optimized tandem (td) IRFP ordered from Twist Biosciences as overlapping gene fragments with the sequences:

ATGGCTGAAGGCAGCGTGGCCCGACAGCCAGACCTTTTGACTTGTGACGATGAACCAATCCACA TACCGGGGGCAATACAACCTCATGGTCTCCTTCTGGCGCTTGCTGCCGACATGACTATAGTGGC CGGCTCTGACAACTTGCCGGAATTGACCGGACTTGCTATTGGGGCGTTGATTGGGCGCTCTGCC GCTGATGTATTTGATTCCGAGACACATAATAGGCTTACTATAGCCCTCGCCGAACCAGGGGCTG CCGTCGGCGCTCCTATAACAGTTGGGTTCACGATGCGAAAAGATGCTGGGTTCATTGGTAGCTG GCATCGCCACGATCAACTTATCTTCCTTGAGCTTGAACCCCCTCAACGGGACGTTGCGGAACCC CAAGCTTTCTTTAGAAGGACCAATTCAGCCATAAGGCGCCTTCAGGCCGCAGAGACATTGGAGT CCGCGTGTGCGGCAGCAGCGCAGGAAGTACGAAAGATCACGGGATTTGACCGGGTTATGATTT ACAGATTCGCATCTGATTTCTCCGGGGAAGTCATCGCGGAGGATCGGTGTGCAGAAGTGGAAA GCAAGCTTGGTTTGCATTACCCCGCATCTACGGTTCCGGCCCAAGCGAGGAGACTGTATACGAT AAACCCAGTGAGGATCATACCTGACATAAATTATAGACCGGTTCCCGTTACGCCAGACCTGAAC CCCGTCACAGGCAGGCCAATAGACTTGTCTTTTGCAATCCTGCGGTCAGTCTCACCTGTTCACCT CGAGTTTATGAGGAACATAGGGATGCATGGGACGATGAGCATCTCAATCCTGAGAGGTGAACGG CTCTGGGGACTTATTGTTTGTCATCATCGCACACCGTATTACGTTGACCTTGATGGTCGCCAGGC CTGCGAACTCGTAGCTCAAGTATTGGCCTGGCAGATCGGTGTTATGGAGGAAAGCGGTCATGG GACTGGGAGTACAGGTAGCGGCAGCTCTAGTGGCACCTCC

and

TAGCGGCAGCTCTAGTGGCACCTCCATGGCAGAAGGGTCCGTAGCAAGGCAACCTGACTTGTT GACCTGTGATGATGAACCGATTCACATTCCTGGAGCAATTCAACCGCATGGGCTGCTCCTTGCTT TGGCAGCGGACATGACGATCGTCGCCGGCTCCGATAACCTGCCCGAGTTGACGGGCTTGGCGA TAGGAGCCCTGATAGGCCGCTCAGCCGCTGACGTATTCGATAGCGAAACGCATAACCGGCTTAC AATCGCCTTGGCTGAACCGGGCGCGGCCGTGGGAGCACCGATTACTGTAGGCTTTACAATGAG AAAAGACGCCGGCTTTATCGGGTCATGGCACCGACATGACCAGCTGATTTTCCTGGAATTGGAG CCCCCGCAGCGGGATGTAGCCGAACCACAGGCCTTCTTCCGGCGCACTAACTCCGCAATTAGG AGACTGCAGGCAGCTGAGACTTTGGAATCAGCATGCGCGGCAGCTGCACAAGAAGTCCGGAAA ATCACGGGTTTTGACCGAGTCATGATCTATAGATTCGCGAGCGATTTCTCAGGAGAAGTTATTGC GGAAGACCGATGCGCGGAGGTAGAATCTAAGCTTGGGTTGCACTACCCCGCCTCCACCGTTCC GGCGCAAGCCAGACGGCTCTATACCATTAATCCGGTGCGGATCATTCCAGATATAAATTACCGG CCTGTACCTGTGACACCGGATTTGAACCCTGTCACGGGCCGACCGATAGACCTCAGCTTCGCTA TATTGCGATCTGTGTCACCGGTCCACCTCGAGTTTATGAGGAATATAGGCATGCATGGTACAATG TCCATTTCCATTCTCCGGGGTGAACGGCTTTGGGGCCTCATCGTTTGTCACCATCGAACACCGT ATTACGTCGATCTCGACGGCAGACAGGCATGTGAGTTGGTCGCTCAGGTACTCGCTTGGCAGAT AGGGGTAATGGAGGAG

PCR fragments were amplified using the following primers:

PiggyBacPuro backbone fwd: ACCTGCGGGGACGTGGAAGAAAACCCGGGTCCAatgaccgagtacaagcccacggtg,

rev: cattccacagggtcgacagtacaagcaaaaag.

8X TOPFlash fwd: cttgtactgtcgaccctgtggaatgaagtgcaggtgccagaacatttctc, rev: GTCGGGCCACGCTGCCTTCAGCCATggtggctttaccaacagtaccgg. tdIRFP1 fwd: ATGGCTGAAGGCAGCGTGGC,

rev: GGAGGTGCCACTAGAGCTGC. tdIRFP2 fwd: TAGCGGCAGCTCTAGTGGCAC,

rev: GGTTTTCTTCCACGTCCCCGCAGGTCAACAAACTTCCGCGACCTTCTCCGCTCCCCTCCTCCATTACCCCTATCTGCCAAGCG.

All above constructs were transformed into Top10 competent cells prepared using Mix & Go E.coli Transformation Kit and Buffer set (Zymo Research #T3002), cultured on LB agar plates to select for antibiotic resistance using standard workflows for molecular cloning and DNA production ^46^. Plasmid DNA was purified using the Zyppy Plasmid Miniprep kit (Zymo Research #D0436). In addition to antibiotic selection, constructs were verified via Sanger sequencing using primers targeting fusion junctions of relevant construct domains.

### Lentiviral Production and Transduction

Production of lentivirus carrying opto-GSK3 was accomplished via co-transfection of pLV_Cry2-tdeGFP-GSK3b, pCMV dR8.91 (obtained from Jared Toettcher’s Lab at Princeton University) and pMD 2.G at a 1: 0.88 : 0.11 *mass* ratio using standard PEI-based transfection procedures^47^ Cells were incubated for 24 hours before replacing with fresh media and allowing for lentiviral production for an additional 48 hours. Supernatant was harvested, filtered through 0.22um filter and added to plated cells for transduction. *NOTE: All steps for lentiviral production, transduction and subsequent maintenance of cell lines were carried out in the presence of far-red light or the complete absence of light in attempt to eliminate the possibility of Cry-2 opto-GSK3 clustering interference with cell growth or virus production.

### Construction of CRISPR gRNA Constructs and Homology-Directed Repair Templates

Genomic edits in 293Ts were carried out in cells constitutively expressing Cas9 to maximize editing efficiency.

#### pCAB_minimal gRNA backbone

A vector expressing guide RNA and Cas9 obtained from Max Wilson was subcloned to remove the unnecessary Cas9 ORF via PCR using the following primers:

fwd: acgcgccctgtagcg

rev: cttaatgcgccgctacagggcgcgtggtacctctagagccatttgtctgc,

assembled, cloned, purified and verified as described in the previous section. The baseline pCab_minimal construct was then subsequently used for production of gRNAs targeting exon1 of the human genomic loci of CTNNB1, CSNK1a1, GSK3B. Primers creating 1-3 (depending on PAM site availability/predicted on/off-target editing scores) unique protospacers targeting the 50-bp window surrounding the 1st codon of each gene were annealed and cloned into the pCAB_minimal via BbsI digestion and ligation (New England BioLabs # R3539S, Takara #6023) using standard protocols ^48^. The following primers were used for sticky-end ligation of protospacers:

CTNNB1_1 fwd: caccgTGAGTAGCCATTGTCCACGC rev: aaacGCGTGGACAATGGCTACTCA

CTNNB1_2 fwd: caccgTGAAAATCCAGCGTGGACAA rev: aaacTTGTCCACGCTGGATTTTCAc

CTNNB1_3 fwd: caccGCGTGGACAATGGCTACTCA rev: aaacTGAGTAGCCATTGTCCACGC

CSNK1a1_1 fwd: caccGGCCAAGCCCCGACACCTCT rev: aaacAGAGGTGTCGGGGCTTGGCC

CSNK1a1_2 fwd: caccgAGGCTGAATTCATTGTCGGA rev: aaacTCCGACAATGAATTCAGCCT

GSK3B fwd: caccCGAAGAGAGTGATCATGTCA rev: aaacTGACATGATCACTCTCTTCG

#### Homology-Directed Repair (HDR) Templates

Blunt-end PCR products were used in conjunction with gRNAs to template genomic edits containing desired knock-ins. Blunt-end, double-stranded HDR templates were created via DNeasy Blood and Tissue genomic prep kit (Qiagen, 69504) of 293T cell line to be edited (see next section) and PCR using primers targeting amplicons of a 500-1000bp window centered on the intended cut site. The following primers were used to amplify genomic loci homology regions:

tdmRuby3: GGcGGTAGTGGcGGcGGAagcATGGTTAGCAAAGGGGAGGAGCTTATAAAGGAAAATATGAGAAT GAAAGTTGTCATGGAAGGTTCAGTGAATGGCCATCAGTTTAAATGTACAGGTGAAGGCGAGGGA CGCCCTTATGAAGGAGTCCAAACTATGAGGATCAAAGTCATAGAGGGAGGTCCTCTCCCCTTCG CCTTCGATATCCTCGCCACCTCTTTCATGTATGGTTCAAGAACATTTATCAAGTATCCTGCCGATA TACCAGACTTCTTTAAGCAGTCATTTCCAGAAGGTTTCACTTGGGAACGAGTCACTAGGTATGAG GACGGCGGGGTTGTGACAGTAACTCAAGACACCTCTTTGGAAGATGGTGAGTTGGTCTACAACG TGAAGGTACGCGGGGTTAATTTCCCTTCTAACGGGCCTGTTATGCAAAAGAAGACAAAGGGTTG GGAGCCAAATACCGAGATGATGTATCCTGCAGATGGTGGCCTGCGGGGCTATACCGACATCGC TCTGAAGGTAGACGGCGGGGGCCACCTCCATTGTAATTTTGTAACCACTTACAGGTCTAAGAAG ACCGTGGGTAACATTAAGATGCCAGGGGTTCATGCTGTCGACCATAGATTGGAGCGGATAGAAG AAAGCGACAACGAGACCTACGTCGTGCAACGCGAAGTCGCAGTAGCCAAGTATTCCAATCTCGG GGGAGGTATGGATGAACTCTATAAAGGCGGATCCGGTGGTGTGTCCAAGGGAGAAGAACTGAT CAAAGAGAACATGAGGATGAAGGTCGTGATGGAGGGCAGCGTCAACGGACACCAATTCAAGTG CACCGGAGAGGGAGAAGGCAGACCATACGAGGGCGTGCAGACAATGAGAATTAAGGTGATCGA AGGCGGACCACTGCCTTTTGCTTTCGACATTCTGGCTACAAGCTTCATGTACGGCAGCAGGACC TTCATTAAATACCCCGCTGACATCCCTGATTTTTTCAAACAAAGCTTCCCTGAGGGCTTTACCTGG GAGAGAGTGACAAGATACGAAGACGGAGGCGTCGTCACCGTCACACAGGATACAAGCCTGGAG GACGGAGAACTGGTGTATAACGTCAAAGTCAGAGGAGTGAACTTTCCCAGCAATGGCCCCGTGA TGCAGAAAAAGACCAAAGGCTGGGAACCTAACACAGAAATGATGTACCCAGCCGACGGAGGAC TGAGAGGATACACAGACATTGCCCTCAAAGTGGATGGAGGAGGACATCTGCACTGCAACTTCGT CACAACCTACAGATCCAAGAAAACAGTCGGAAATATCAAGATGCCTGGCGTGCACGCCGTGGAT CACAGGCTGGAAAGGATTGAGGAGTCCGATAATGAAACATATGTGGTCCAGAGGGAGGTGGCC GTCGCTAAATACAGCAACCTGGGCGGCGGCATGGACGAGCTGTACAAGGGGGGATCAGGAGGa GGctct

CTNNB1 fwd: ATAAAAAGACATTTTTGGTAAGGAGGAGTTTTCACTGAAGTTCAGCAGTGATGGAGCTGTGGTTG AGGTGTCTGGAGGAGACCATGAGGTCTGCGTTTCA CTAACCTGGTAAAAGAGGATATGGGTTTTTTTTGTGGGTGTAATAGTGACATTTAACAGGTATCC CAGTGACTTAGGAGTATTAATCAAGCTAAATTTAAATCCTAATGACTTTTGATTAACTTTTTTTAGG GTATTTGAAGTATACCATACAACTGTTTTGAAAATCCAGCGTGGACAGGcGGTAGTGGcGGcGGA

agc

rev: TAGGGAACCACCTAACAGTTACTCACTGAATCAGTGGAAGAATGGTACTGCATCCAGGCTCCAG AAGCAGTCATCCAGACTAGATTCCTGCTGGTGGCTT GTTTGCTATTTCACCAAGCCATTAGGAGGAGTGAGCAGAAAATGGAGCAAAAGGTAGCCTGACA AGTAAGCAGGGAGAGAGGAAAGCAGGGGGATCTCAGCCAGACTGGCTTAATGGCAACGAAGCA GAGCCCCAATTCAGTAACTAAAGATTTAATGACACAAACCTTGAGTAGCCATagagCCtCCTCCTG ATCCCCC

*NOTE: CTNNB1 homology arms were synthesized (requiring no genomic amplification step) and provided as a generous gift from Integrated DNA Technologies.

CSNK1a1fwd: CCAGCCCGCGACGTC rev: CTTGACCCTTTTAGGGAGACAGCG

GSK3B fwd: GATTTGCCCTCTCTTTTCTCTCCTCC rev: CCAAATAAATATCATATTATCTCAATTCAAGGTTAATGAGACCG

The above amplicons were then used in a second round of PCR to obtain separate upstream and downstream homology arms that flanked desired knock-ins and overlap extension was used to construct the final desired amplicons bearing tdmRuby3 and 7AA GS linker. The following primers were used:

Generic tdmRuby3 insert fwd: GGcGGTAGTGGcGGcGGAagcATGGTTAGCAAAGGGGAGGAGC, rev: agagCCtCCTCCTGATCCCCCCTTGTACAGCTCGTCCATGCC

CSNK1a1 upstream homology arm rev: gctTCCgCCgCCACTACCgCCCCTGAGAGACGAAGATGGAGGC

CSNK1a1 downstream homology arm fwd: GGGGGATCAGGAGGaGGctctATGGCGAGTAGCAGCGGC

GSK3B upstream homology arm rev: gctTCCgCCgCCACTACCgCCGATCACTCTCTTCGCGAATCACC

GSK3B downstream homology arm fwd: GGGGGATCAGGAGGaGGctctATGTCAGGGCGGCCC

*NOTE: Original upstream fwd and downstream rev primers listed above for isolating genomic loci were reused in the present step and thus not repeated here.

### CRISPR-Cas9 Fluorescent Tagging

Bare 293T cells were first co-transfected using PEI^47^ with the H2B-mTagBFP2 vector and Super PiggyBac Transposase-expressing vector (System Biosciences Inc. # PB210PA-1) via polyethylenimine (Sigma #408727-100mL) transfection reagent and standard workflows ^47^. Cells were allowed 72 hours following transfection to reach steady-state expression of integrated construct and were enriched via 2 rounds of fluorescence-activated cell sorting (FACS, SH800S, Sony Biotechnology) for cells fluorescent in the 450nm excitation (blue) channel: a bulk enrichment to obtain a largely ‘positive’ population and a 2^nd^ to obtain clonal populations. A high-expressing clone was expanded and used as a ‘chassis’ cell line for subsequent CRISPR editing.

CRISPR chassis cells were then co-transfected with one of the constructed gRNA plasmids and respective HDR templates at a 2:1 HDR template:gRNA plasmid molar ratio and allowed 72 hours to reach steady-state expression. Similar to the process described above, cells were subject to 2 rounds of FACS (561nm excitation, red laser) to obtain a clonal population. Knock-in validation was accomplished via a combination of fluorescence microscopy, genomic PCR and sequencing (using primers for initial amplification of loci and construction of HDR templates). In the case of all intended knock-ins, spatiotemporal fluorescence expression of cell populations was binary (either fluorescent or not) and uniform (no detected variation in brightness or localization between fluorescent clones), suggesting that selected clones were broadly representative of overall edited populations.

### Development of Inducible Axin1, APC and β-catenin Cell Lines

293Ts were co-transfected as described in the previous section with PiggyBac and compatible XLone-Axin-tdmRuby3 and pPig_CuO-APC-tdmIRFP670::CymR expression cassettes. 72 hours following transfection cells were selected in 1uM Blasticidin (Invivogen, #ant-bl-05) and 100ug/mL Hygromycin B Gold (Invivogen, #ant-hg-1). Blast+/Hygro+ cells were then clonally sorted via FACS as described in the previous section to obtain a uniform population for experiments. For iPSCs, Both Piggyback and Donor plasmids were chemically transfected when cells reached 30% confluency using Lipofectamine™ Stem Transfection Reagent (manufactures protocol). Following transfection Blasticidin selection (1uM) was initiated 5 days later. At the end of Blasticidin selection, 12 clones were manually picked under a dissection microscope and continuously cultured in Blasticidin (1uM) for an additional week. Upon fluorescence signal confirming successful integration, Blasticidin (1uM) treatment ceased and 1 clone was chosen for the remaining experiments.

### Small Molecules

CHIR 99021 (STEMCELL TECHNOLOGIES # 72052) was resuspended in dimethyl sulfoxide according to supplied manufacturer recommendations and diluted to 5X concentrated stocks in culture medium immediately prior to use on cells. In all cases CHIR was used at 10uM. Doxycycline hyclate (Sigma Aldrich # D9891-1G) was resuspended in phosphate-buffered saline and diluted to 5X desired concentration in culture medium prior to use. Stock cumate solution (System Biosciences # QM100A-1) was diluted to 5X in culture medium prior to use.

“Low” dose of Dox referred to in **Fig. 2C**,**D** in the context of Axin and APC induction was 20ng/mL concentration in culture medium, “High” dose was 200ng/mL. “Low” dose of Cumate was 100ng/mL, “High” was 1mg/mL. The dose of Dox used in β-cat induction in **Supp. Fig. 4I**,**J** was 100ng/mL.

### Wnt-3a treatments

Recombinant Human Wnt-3a (R&D Systems 5036-WN-010) was resuspended in in PBS containing 0.1% BSA according to supplied manufacturer recommendations and diluted to 5X concentration in culture medium immediately prior to use. In all cases Wnt-3a was used at a final concentration of 1ug/mL.

### Antibodies, Immunofluorescence and Western Blot

Primary antibodies used for immunofluorescent markers of the centrosome were α-GM130 (BD 610822, 1:1000 dil.) and α-γ-tubulin (Sigma Aldrich T5326-25UL, 1:1000). Secondary used for both stains was α-Ms-Alexa-488 (Invitrogen A28175, 1:1000). Tissue fixation and staining was carried out using standard protocols using cold methanol^49^ Immunofluorescent samples were imaged using confocal microscopy (see below). Antibodies used for Western Blotting and immunofluorescence were α-β-catenin (Cell Signaling, # 2698S, 1:1000) and α-β-actin (Sigma, A3853, 1:1000). Secondary antibodies used were α-Gt-680RD and α-Ms-800CW (Licor 926-6807 and 926-32212 respectively, both 1:10,000 dil.). Standard immunoblot procedures were used^50^.

### Imaging

All live and fixed cell imaging experiments were carried out using a Nikon W2 SoRa spinning-disk confocal microscope equipped with incubation chamber maintaining cells at 37°C and 5% CO2. Glass-bottom culture plates (Cellvis # P96-1.5H-N) were pre-treated with bovine fibronectin (Sigma #F1141) in the case of 293Ts or Matrigel in the case of H9 and iPSCs, and cells were allowed to adhere to the plate before subsequent treatment or imaging. Fluorescence recovery after photobleaching was performed via custom Nikon NIS Elements JOBs function and 488nm FRAP laser (Nikon LUN-F laser unit, 100mW power output from the APC fiber tip).

### Optogenetic Stimulation

Spatial patterning of light during timelapse fluorescent imaging sessions was accomplished via purpose-built microscope-mounted LED-coupled digital micromirror devices (DMDs) triggered via Nikon NIS Elements software. Stimulation parameters (brightness levels, duration, pulse frequency) were optimized to minimize phototoxicity while maintaining continuous activation of Cry-2. For DMD-based stimulation on the microscope, the final settings for ‘Light ON’ were 25% LED power (λ = 455nm), 2s duration pulses every 30s. For experiments that did not require frequent confocal imaging, cells were stimulated via a benchtop LED array purpose-built for light delivery to cells in standard tissue culture plates (‘OptoPlate’) adapted from previously established designs^51^. The same light delivery parameters were used for OptoPlate based stimulation as for microscope mounted DMDs. Light was patterned to cover the entire surface of intended wells of plates used, rather than a single microscope imaging field.

### Image Analysis

All quantification of raw microscopy images was carried out using the same general workflow: background subtraction > classification > measurement > normalization > statistical comparison. Subcellular segmentation of nuclear fluorescence was performed via custom Matlab scripts using H2B-mTagBFP2 brightness, size and circularity to mask objects. When experimental conditions did not permit segmentation via H2B-mTagBFP2 nuclear fluorescence (such as with live-cell optogenetic stimulation) cells were selected at random using custom ImageJ macro that generates random ROIs (available upon request). Unless otherwise noted, mean fluorescent intensity of regions of interest were measured and subsequently processed. Raw measurements were compiled, processed, and plotted via custom Matlab scripts, available upon request.

### Statistical Analysis

#### Significance key

| p value range | Symbol |
| --- | --- |
| $0.01 < p < 0.05$ | * |
| $0.001 < p < 0.01$ | ** |
| $p < 0.001$ | *** |

All statistical tests were carried out on final grouped datapoints presented in figures using independent samples t-tests (Matlab function “ttest2”) except for **Supp. Fig. 4D** which was the result of one-way ANOVAs.

### Simulation Methods

We used the python-based FEniCS computing environment (https://fenicsproject.org/) to solve the modified Cahn-Hilliard partial differential equations using the Finite Element Method (FEM). The Cahn-Hilliard equation, in its general form, is a parabolic equation with first-order time derivatives, and second- and fourth-order spatial derivatives. To solve this equation using a standard Lagrange finite element basis the equation is recast as two coupled second-order equations:

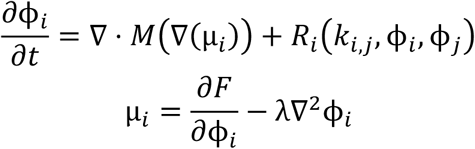

Where the free energy function F is defined as

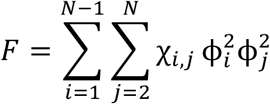

and R_i_ is the added reaction term with such that

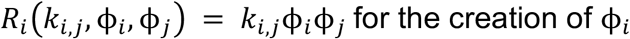

and

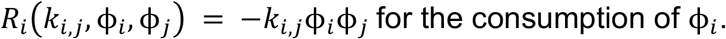

The system is time discretized according to established methods ^52^. Assuming that the total free energy of the system decreases to a minimum with time, we use the built-in Newtonian solver in the FEniCS environment to approximate the forward evolution of the system in time. To represent the enzyme activities in the DC we model only clients, with scaffold existing implicitly as the interaction parameters between system components. Representations are:

| Component Variable | Component Name | CH Equation |
| --- | --- | --- |
| $\phi_1$ | GSK1β | $\frac{\partial \phi_1}{\partial t} = M_1 \nabla^2 \mu_1$ |
| $\phi_2$ | CK1 | $\frac{\partial \phi_2}{\partial t} = M_2 \nabla^2 \mu_2$ |
| $\phi_3$ | β-Catenin | $\frac{\partial \phi_3}{\partial t} = M_3 \nabla^2 \mu_3 - (k_{2,3} \cdot \phi_2 \cdot \phi_3)$ |
| $\phi_4$ | Phospho-S45 β-Catenin | $\frac{\partial \phi_4}{\partial t} = M_4 \nabla^2 \mu_4 + (k_{2,3} \cdot \phi_2 \cdot \phi_3) - (k_{1,4} \cdot \phi_1 \cdot \phi_4)$ |
| $\phi_5$ | Phospho-S33/S37/S45/T41 β-Catenin | $\frac{\partial \phi_5}{\partial t} = M_5 \nabla^2 \mu_5 + (k_{1,4} \cdot \phi_1 \cdot \phi_4)$ |
| $\phi_6$ | Cytoplasm | $\frac{\partial \phi_6}{\partial t} = M_i \nabla^2 \mu_i$ |
| $\phi_7$ | Nucleator | $\frac{\partial \phi_7}{\partial t} = M_i \nabla^2 \mu_i$ |

### Interaction Parameter

One of the key factors that tunes system behavior is the interaction parameter χ. Assuming a system with constant temperature and pressure, the interaction parameter determines the free energy of the system. When χ is positive between two components, the system will tend to de-mix. If χ is negative between two components they will tend to mix. Lastly, if χ is neutral, the two components are interaction-less. For simplicity, we limited interactions to one of three types: binding (χ ∼ -0.1), neutral (χ ∼ 0), and separating (χ ∼ 2). As noted above we represent the binding action of DC scaffolds implicitly. Scaffold interactions are taken to be of similar strength and were obtained from literature values described in the table below:

| Interaction | Behavior | Source |
| --- | --- | --- |
| scaffold to GSK3β | Binding | Lee 2003, Pronobis 2017 |
| scaffold to CK1 | Binding | Lee 2003, Pronobis 2017 |
| scaffold to β-cat | Binding | Lee 2003, |
| scaffold to P1 β-cat | Binding | Schaefer 2019 |
| scaffold to P4 β-cat | Neutral | Lee 2003, Schaefer 2019 |
| scaffold to cytoplasm | Separating | This study |
| scaffold to centrosome | Binding | This study, Fumoto 2009 |

Given that the APC/Axin interacts with the DC proteins, the following interaction constants were selected for the system with implicit Axin. We set mixing = 2.0, neutral = 0.0, and de-mixing = -0.1. The following table shows the selected interaction behavior

### Simulation Flow

First all parameters are defined (χ, λ, dt, and M). We generate a grid mesh with a closed boundary conditions to mimic the closed system within a cell. A layer is generated for each simulated component and +-5% noise of the initial value is added to induce inhomogeneities. The FEniCS package partial differential solver is called to generate the chemical potential with respect to each component. The final step is to define the output file path and then use the built-in newton solver to generate the simulation. The simulations are then rendered using Paraview software. A detailed python notebook of the simulations is available on https://github.com/MZWLab/Lach2022.

## Supplemental Figures

**Supp. Fig. 1:**
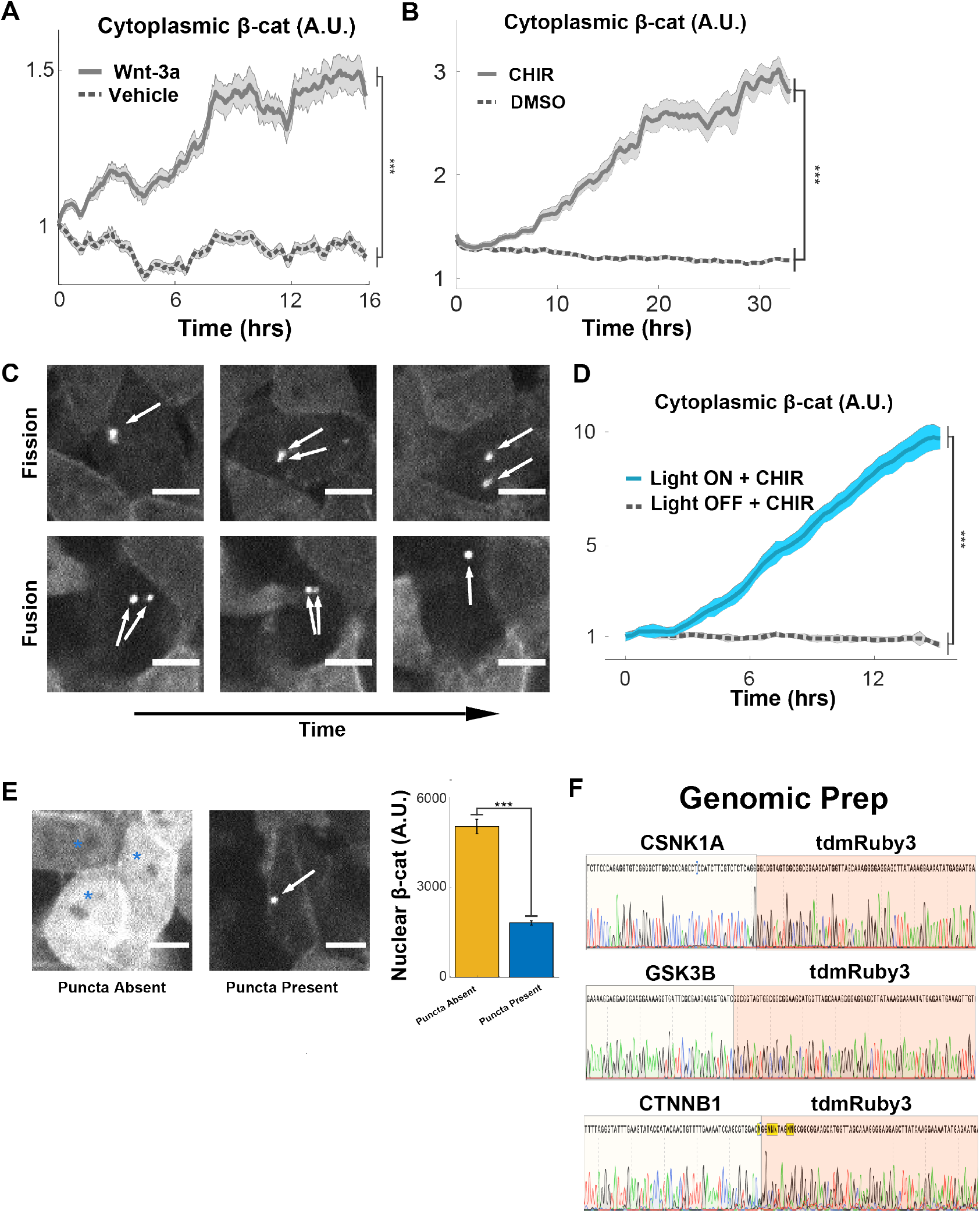
Endogenously expressed β-catenin puncta are inversely correlated with CHIR-mediated Wnt pathway activation and β-catenin accumulation and show hallmarks of dynamic liquidity. **A-B**. Measurements of CRISPR cytoplasmic tdmRuby3-β-catenin in live 293Ts, data presented as mean fluorescent intensity fraction of t0 +/- s.e.m. (N = 30 cells per condition). **C**. Montage of single CHIR+ cells containing β-catenin puncta undergoing fission and fusion over time. **D**. Measurements of CRISPR cytoplasmic tdmRuby3-β-catenin in live 293Ts treated with CHIR, with or without blue light stimulation, data presented as mean fluorescent intensity fraction of t0 +/- s.e.m. (N = 30 cells per condition). **E**. *Left:* Representative images of tdmRuby3-β-catenin cells +CHIR for 24hrs. Arrows indicate puncta, asterisks indicate puncta absent. *Right:* Comparison of mean nuclear β-catenin fluorescence between +CHIR cells with and without visible β-catenin puncta. **F**. Sanger sequencing traces from genomic PCRs targeting 5’ endogenous loci of CRISPR tdmRuby3 knock-ins. Red regions indicate tdmRuby3 insert.

**Supp. Fig. 2:**
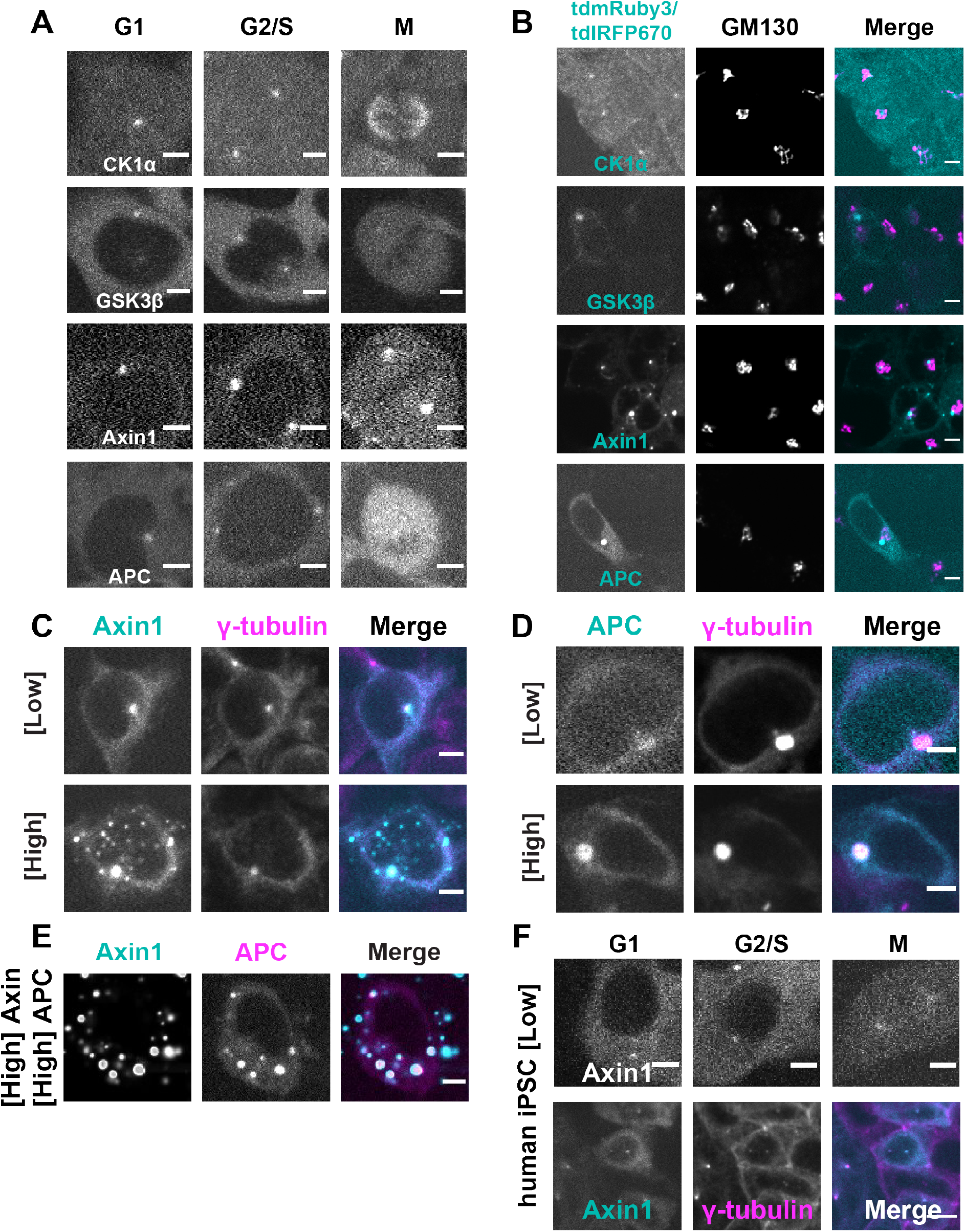
Centrosomal Destruction Complex droplets are spatially correlated with cell cycle progression. **A**. Representative images of indicated Destruction Complex (DC) components taken live, at various stages of the cell cycle. Montages follow the same cell through time. Scale = 10μm **B**. Representative fixed images of indicated DC components stained for endogenous GM130. Scale = 10μm. **C**. Representative images of fixed cells with varying Axin1 induction and co-stained for endogenous γ- tubulin. **D**. Representative images of fixed cells with varying APC induction and co-stained for endogenous γ-tubulin. **E**. Representative images of live cells induced to high Axin1 and APC levels simultaneously. **F**. *Upper:* Representative images of live human induced-pluripotent stem cells (iPSCs) at various stages of the cell cycle. *Lower:* Images of fixed human iPSCs co-stained for endogenous γ- tubulin.

**Supp. Fig. 3:**
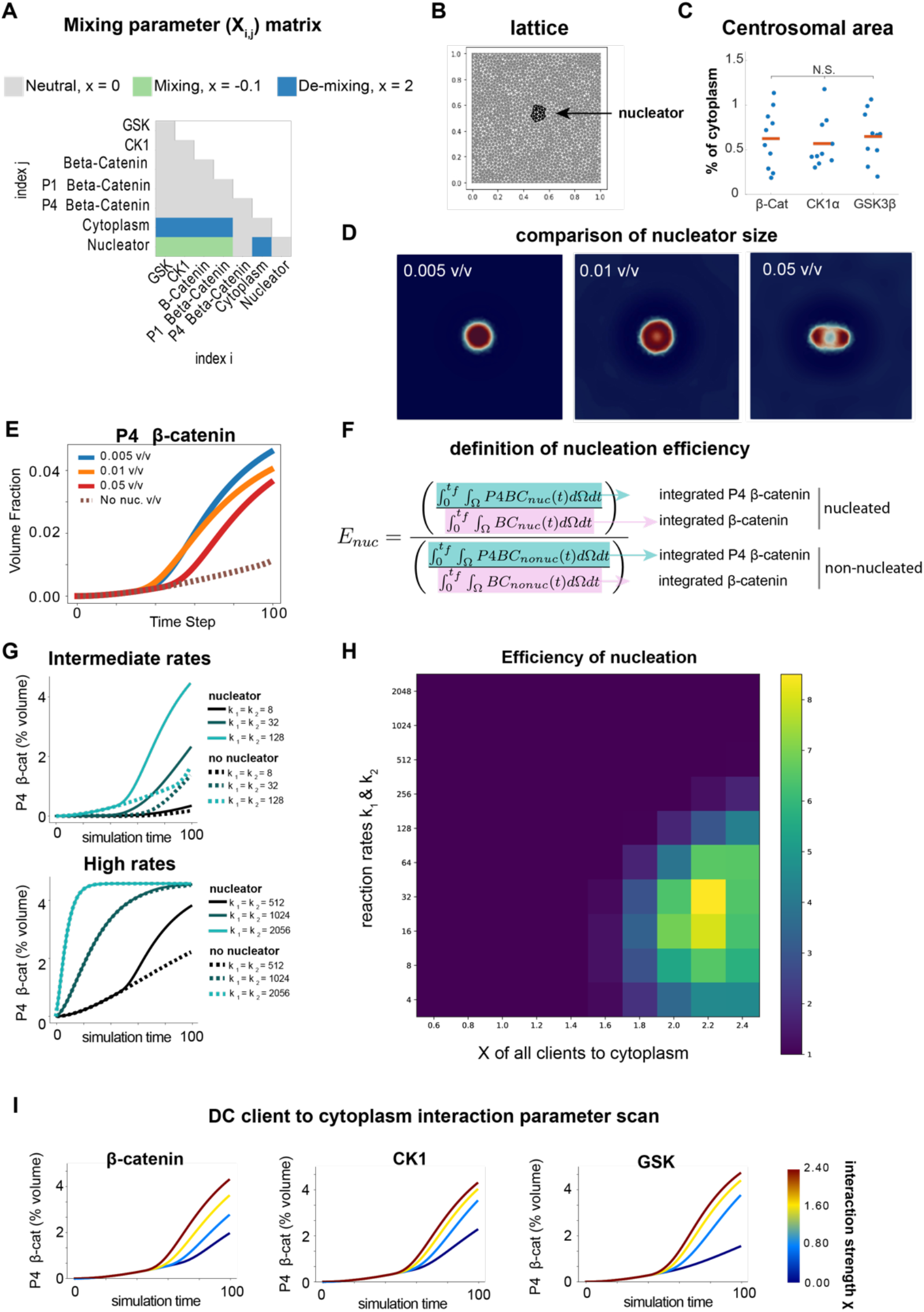
Exploring In Silico Model of Centrosome driven Phase Separation. **A**. Interaction matrix of each component in the model. Gray indicates a neutral state, blue represents de-mixing, and green represents mixing **B**. Example lattice used to model the system with example nucleator region in black. After initial conditions are assigned, a model of diffusion operates on grid positions based on modified Cahn Hilliard equations. **C**. Quantification of the area of centrosomal droplets in comparison of total cell volume taken from CRISPR-tagged cells. Mean is represented by red line. **D**. Demonstration of the effects of nucleator size on system nucleation process. With a smaller centrosome, the droplet is more densely packed with enzymes whereas a larger centrosome results in droplet separation. **E**. Quantification of the effect of centrosome size on P4 β-catenin generation. **F**. Definition of nucleation efficiency as the ratio of the quotient of P4 β -catenin and β-catenin in a nucleated versus an unnucleated system. **G**. P4 β-catenin accumulation in log2 scan of kinase reaction rates. **H**. Nucleation efficiency of as a function of reaction rates and X (interaction parameter) of all clients and the cytoplasm. **I**. Quantification of in silico models of “opto”-β-catenin, “opto”-CK1, and “opto”-GSK. The graphs show increased gain from “opto”-GSK driven separation.

**Supp. Fig. 4:**
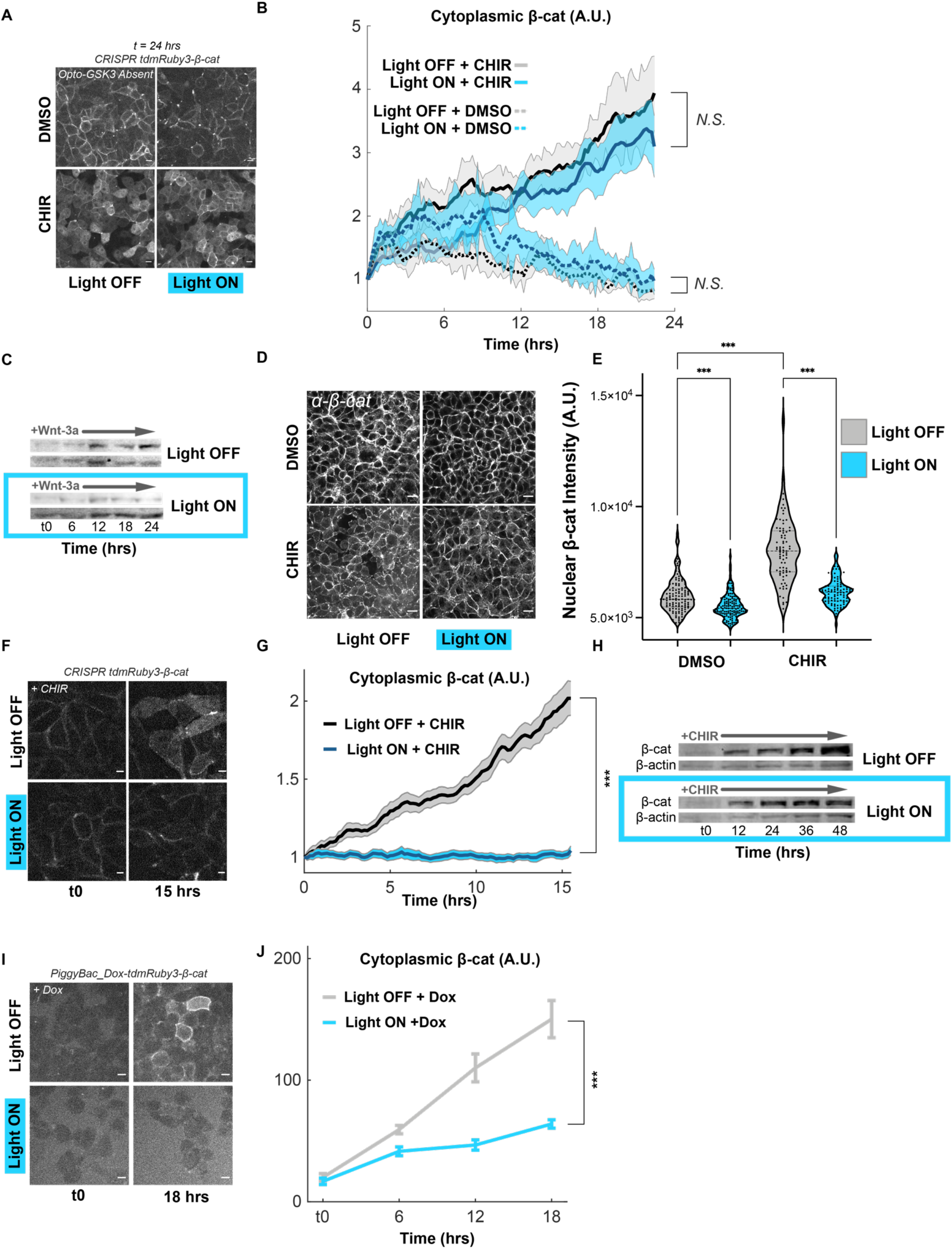
Opto-GSK3 suppresses β-catenin accumulation due to GSK3β inhibition or exogenous chemical induction. **A**. Representative images of live cells treated with CHIR or DMSO vehicle, with or without blue light stimulation. Scale = 10μm. **B**. Measurements of experiment shown in **A**. Data presented as mean +/- s.e.m. (N=30 cells per condition). **C**. Representative Western blots of lysates from 293Ts bearing Opto-GSK3 and treated with Wnt-3a, with or without blue light stimulation for the indicated time course. **D**. Representative images of cells bearing Opto-GSK3 fixed and stained for endogenous β-catenin after culture in the indicated conditions for 48 hrs. **E**. Violin plots of cells from **D. F**. Representative images of 293Ts bearing Opto-GSK3 and endogenously-expressed tdmRuby3-β-catenin treated with CHIR, with or without blue light stimulation for 24hrs. Measurements from experiment shown in **G:** lines represent fold-change from t0 means +/- s.e.m. for cells in each condition (Light ON N=67, Light OFF N=50 cells). **H**. Representative Western blots of lysates from 293Ts bearing Opto-GSK3 and treated with CHIR, with or without blue light stimulation for the indicated time course. **I**. Representative images of 293Ts bearing Opto-GSK3 and Dox-inducible -β-catenin-tdmRuby3 treated with Dox, with or without blue light stimulation. Scale = 10μm. **J**. Quantification of experiment in **I:** lines represent absolute means +/- s.e.m for cells in each condition (Light ON N=28, Light OFF N=27 cells).

