## Supplementary Video Captions for "Nucleation of the destruction complex on the centrosome accelerates degradation of β-catenin and regulates Wnt signal transmission"

**Supp. Vid. 1: Cells with β-cat Puncta Resist β-cat Accumulation in response to CHIR.** *Left:* Imaging fields of live tdmRuby3-β-cat cells treated with DMSO control. *Right:* Imaging fields of tdmRuby3-β-cat cells treated with CHIR. Arrows indicate β-cat puncta.

**Supp. Vid. 2: Activation of Cry-2-Lrp6c Induces β-cat Accumulation** *Left*: Imaging fields of live, unstimulated tdmRuby3-β-cat, Cry2-Lrp6c cells. *Right:* Imaging fields of tdmRuby3-β-cat, Cry2-Lrp6c cells stimulated with blue light throughout indicated timecourse. Videos were taken from cells in the same well.

**Supp. Vid. 3: Activation of Cry-2-Lrp6c Results in Dissolution of β-cat Puncta** Zoomed videos of cells presented in **Supp. Vid. 2.** Arrows indicate β-cat puncta.

**Sup Video 4: In-silico behavior of destruction components with a centrosomal region.** In-silico model of phase separation behavior for every component involved in a hypothetical WNT pathway over 100 simulation time steps in the presence of a centrosome.

**Sup Video 5:** **In silico behavior of destruction components without a centrosomal region.** In-silico model of phase separation behavior for every component involved in a hypothetical WNT pathway over 100 simulation time steps without the presence of a centrosome.

**Sup Video 6: Impact of interaction parameter χ on destruction complex component behavior.** In-silico model of the destruction components (CK1α, GSK3β, and β-catenin) at various interaction parameter values (χ) over 100 simulation time steps showing that increasing χ increases separation propensity.

**Supp Vid 7: Activation of Opto-GSK3 Increases Centrosomal Condensate Partitioning.** Zoomed video of Opto-GSK3 cells stimulated with blue light throughout indicated timecourse.
